# Characterization of *Arabidopsis* aldolases AtFBA4 and AtFBA5; inhibition by morin and interaction with calmodulin

**DOI:** 10.1101/2024.03.29.587371

**Authors:** Kyle Symonds, Milena A. Smith, Oona Esme, William C. Plaxton, Wayne A. Snedden

**Affiliations:** Department of Biology, Queen’s University, Kingston, Ontario, Canada, K7L3N6

**Keywords:** Arabidopsis, fructose bisphosphate aldolase, calmodulin, glycolysis, enzyme kinetics

## Abstract

Fructose bisphosphate aldolases (FBAs) catalyze the reversible cleavage of fructose 1,6-bisphosphate into dihydroxyacetone phosphate and glyceraldehyde 3-phosphate. We analyzed two previously uncharacterized cytosolic *Arabidopsis* FBAs, AtFBA4 and AtFBA5. Based on a recent report, we examined the interaction of AtFBA4 with calmodulin (CaM)-like protein 11 (AtCML11). AtFBA4 did not bind AtCML11, however, we found that CaM bound AtFBA5 in a Ca^2+^-dependent manner with high specificity and affinity (*K_D_* ∼ 190 nM) and enhanced its stability. AtFBA4 and AtFBA5 exhibited Michaelis-Menten kinetics with *K*_m_ and *V*_max_ values of 180 µM and 4.9 U/mg for AtFBA4, and 6.0 µM and 0.30 U/mg for AtFBA5, respectively. The flavonoid morin inhibited both isozymes. Our study suggests that Ca^2+^ signalling and flavanols may influence plant glycolysis/gluconeogenesis.

## Introduction

Fructose bisphosphate aldolases (FBAs, EC 4.1.2.13) are ubiquitous across taxa where they play pivotal roles in glycolysis and gluconeogenesis by catalyzing the reversible aldol cleavage of fructose-1,6-bisphosphate (FBP) into dihydroxyacetone phosphate (DHAP) and glyceraldehyde-3-phosphate (G3P). Based on their different catalytic mechanisms, FBAs can be divided into two classes: class I, metal-independent, and class II, metal-dependent enzymes. Class II FBAs, require divalent metal cation cofactors such as Zn^2+^ and are predominantly found in microorganisms, whereas class I FBAs occur in eukaryotes and use a Lys residue in the active site as a Schiff base for the aldol reaction intermediate (1). Intriguingly, despite their low sequence identity, class I and II FBAs exhibit similar secondary and tertiary structures. While FBAs typically form homotetramers, a subclass of FBAs, class Ia, is an archaeal ortholog known to assemble into octamers, decamers, or even higher-order complexes despite sharing the same reaction mechanism of class I FBAs (2).

The Arabidopsis genome encodes eight FBA isozymes, three of which (FBAs1,2,3) localize to the plastids and the remaining five to the cytosol (3). Arabidopsis mutants lacking chloroplastic FBA isozyme AtFBA2 or AtFBA3 display reduced growth, and loss of both AtFBA1 and AtFBA2 is lethal (4). However, the enzymatic properties of most plant FBA isoforms remain unclear. For example, in Arabidopsis, only AtFBA6 and AtFBA8 have been characterized biochemically (5, 6). Similar to other glycolytic enzymes, FBAs are thought to possess additional moonlighting functions in plants (1, 32). Garagounis et al. (2017) demonstrated the ability of FBA8 to bind F-actin filaments but not G-actin, suggesting a potential role for FBAs as actin-bundling proteins *in vivo*. This finding aligns with previous reports in animals that showed the interaction of FBAs with actin filaments and their contribution to micro-compartmentation in cells (7).

A study using *Medicago sativa* and *Medicago truncatula* highlighted the increased freezing resistance that resulted from elevated FBA gluconeogenic activity under cold stress (8). This phenomenon was reported to be a consequence of the interaction of *M. sativa* calmodulin-like 10 (MsCML10) with MsFBA6, leading to enhanced gluconeogenesis and the accumulation of sugars as osmoprotectants (8). These observations suggest a possible Ca^2+^-dependent regulation of FBA during abiotic stress response, where compatible solutes help plants acclimate to stresses such as cold, heat, salinity, and drought. It is noteworthy that these abiotic stresses utilize Ca^2+^ signalling pathways and Ca^2+^ sensors such as calmodulin (CaM) to propagate information by interacting with various downstream targets (9). CaM targets include transcription factors, channels, pumps, and various other proteins that, collectively, carry out cellular responses to external stimuli (9, 10). A notable example of a CaM-regulated metabolic enzyme in plants is glutamate decarboxylase, whose activity helps to maintain carbon flow into the Krebs cycle during stress (11). In addition to multiple isoforms of the evolutionarily-conserved CaM, plants also possess a large family of CaM-like (CML) proteins that are not found in animals. The genetic model, Arabidopsis, has seven CaM isoforms and 50 CMLs (12, 13), most of which have not been thoroughly studied.

Native plant FBAs have been previously purified and biochemically characterized from several species and tissue types. For example, cytosolic FBAs from carrot storage roots, germinating castor oil seeds, and mung bean were purified to homogeneity and characterized *in vitro* (14–16). Their *K*_m_(FBP) varied between 0.16 µM for castor bean FBA, 6 µM for carrot FBA, and 17 µM for mung bean FBA, whereas corresponding *V_max_* values ranged from ∼2.5 to 26 µmol FBP cleaved/min mg^−1^ (U) (14–16). Several metabolites have been reported as *in vitro* inhibitors of FBA activity, including ATP, phosphoenolpyruvate, and ribose-5-phosphate, likely to coordinate glycolytic flux to meet the metabolic demands of the cell (14–16). Although these inhibitors act on some FBAs, the maximal inhibitory effect and *K*_i_ values reported vary widely, and thus general mechanisms of feedback control of FBAs *in vivo* remain unclear. In addition, AtFBA6, an Arabidopsis cytosolic FBA, is regulated by redox-based, post-translational modifications (5).

As FBAs play a central role in glycolysis, there is interest in them as potential drug targets in parasites. A recent study identified the plant flavonoid morin (3,5,7,2′,4′-pentahydroxyflavone) as a putative class-I FBA inhibitor of EtFBA2 from the protozoan parasite *Eimeria tenella* (17). Morin is derived from the common plant flavonoid quercetin, is primarily found in mulberry plants, and possesses various therapeutic properties in animals, including anti-inflammatory, anti-bacterial, and anti-tumor effects (18). The effect of morin on plant FBAs has not been reported.

Despite their importance in plant central metabolism, FBA isozymes are understudied compared to their orthologs in other organisms. The main objectives of the current study were to biochemically characterize two *Arabidopsis thaliana* cytosolic FBA isozymes that have not been previously examined, AtFBA4 and AtFBA5, and to investigate the binding of AtCML11 to AtFBA4. The latter objective was based on a recent report indicating that orthologs of AtCML11 and AtFBA4 in *M. sativa*, MsCML10 and MsFBA6, respectively, physically interact and impact MsFBA6 activity and the accumulation of compatible solutes during stress (8). Our third objective was to assess whether morin inhibits plant FBA *in vitro* activity as has been reported for a protozoan FBA (17).

## Materials and Methods

### Cloning

*AtFBA4* (At5g03690) and *AtFBA5* (At4g26530) full-length cDNA clones were acquired from Arabidopsis Biological Resource Center and PCR amplified by Q5 DNA Polymerase kit (New England Biolab) before subcloning into the pET28-SUMO or pCAMBIA1300-N-Luc (NLuc) vectors and sequenced via Sanger sequencing (TCAG, SickKids Hospital, Toronto). The pCAMBIA-C-Luc (CLuc) vectors with the cDNAs of *AtCML8, AtCML11, AtCML42,* and *AtCaM7* were described previously (19, 20).

### Expression and Purification

Recombinant SUMO-AtFBA expression was performed in *Escherchia coli* strain BL21 (CPRIL) with the chaperone co-plasmid pACYC (21). Transformed *E. coli* was grown to an OD_600_ of 0.6 at 37°C before inducing expression with a final concentration of 0.5 mM IPTG and expressing for 16-18 h at 16°C. Expression and isolation of His-tagged recombinant proteins, as well as untagged recombinant AtCML11 and AtCaM7 (an evolutionarily conserved isoform), were performed as described (20). Protein purity was analyzed by SDS-PAGE and concentrations determined by the Bradford Assay with BSA as the protein standard. Immunoblotting was performed as reported (20) using anti-His (GenScript) or anti-carrot FBA (14) primary antisera, horse-radish-peroxidase conjugated secondary antiserum (Sigma), and a ChemiDoc^TM^ Touch Imaging System (Bio-Rad).

### Estimation of native molecular mass by gel-filtration FPLC

The native molecular masses of the recombinant FBAs were determined by FPLC as described (22) at a flow rate of 0.2 ml/min on a calibrated Superdex 200 10/300 GL column equilibrated in Tris-buffered saline (TBS; 50 mM Tris-Cl, 150 mM NaCl, pH 7.5) and calculated from a plot of *K_av_* (partition coefficient) against log(molecular mass) using the following protein standards: ferritin (440 kDa), catalase (250 kDa), FBA (158 kDa), BSA (66 kDa), ovalbumin (44 kDa), carbonic anhydrase (29 kDa), chymotrypsinogen A (25 kDa), ribonuclease A (13.7 kDa) and cytochrome c (12.4 kDa).

### FBA Activity Assays

The activities of FBA enzymes were assayed as described (15). Briefly, AtFBA4 or AtFBA5 were diluted to 0.05 mg/mL into TBS containing various concentrations of FBP (Sigma), 0.1 mM NADH (Sigma), 10 U triose-P isomerase and 1 U G3P dehydrogenase (Boehringer Ingelheim) to a final volume of 200 µL per well in a 96-well polystyrene microplate. The activity was measured at 25°C by continuously monitoring NADH utilization at 340 nm using a SpectraMax microplate reader. One unit (U) of activity is defined as the amount of enzyme resulting in cleavage of 1 mol/min. In all cases the rate of reaction was proportional to concentration of enzyme assayed and remained linear with respect to time. For pH-activity determinations, TBS was replaced by 50 mM Bis-tris propane (Sigma) and 50 mM (N-morpholino)ethane sulfonic acid (Sigma) as described (22).

### Thermal Inhibition Assays

Concentrated enzyme stocks (∼1 mg/mL) were aliquoted into PCR tubes with or without CaM7 in molar excess with 1 mM CaCl_2_ or 1 mM EGTA and incubated for 20 min in a thermocycler over a range of temperatures depending on the enzyme’s T_1/2_. Immediately after the incubation period the enzyme solution was aliquoted into a 96-well plate and diluted with the FBA assay mixture and activity monitored as before. Relative activity was plotted against the incubation temperature and the T_1/2_ was calculated from the sigmoidal graph using GraphPad Prism 10.

### Split-luciferase Assay

Split-luciferase (SL) protein-interaction assays were carried out as reported (23). *Agrobacterium tumefaciens* strain GV3101 carrying the plasmids of interest and the p19 silencing suppressor were infiltrated into six-week-old *N. benthamiana* leaves. Whole leaves were removed after 4 days of incubation and were sprayed with a solution containing 1 mM D-luciferin (GoldBio) and 0.01% Silwett L-77 (Lehle Seeds Inc) and left to incubate for 10 minutes in the dark. Leaves were imaged on a ChemiDoc^TM^ Touch Imaging System (Biorad Inc.). Alternatively, leaf discs were collected after 4 days, incubated in 100 µL of water containing 1 mM D-luciferin in a 96-well plate for 10-15 min in darkness, and luminescence was recorded using a SpectraMax Paradigm (Molecular Devices Inc.). Expression of the C-Luciferase fusions was verified by immunoblotting (Santa-Cruz Biotech).

### Calmodulin-Agarose Binding Assay

Tests of AtFBAs binding to CaM-agarose were performed essentially as described (24). Purified recombinant AtFBA isozymes (100 µg) were diluted to 0.5 mg/mL in TBS containing either 1 mM CaCl_2_ or EGTA as denoted in the figures. CaM-linked agarose beads (Agilent) were equilibrated in the same buffer before adding the FBA preparations and incubating for 5 min at room temp. The column was washed with 10 volumes of equilibration buffer and the first wash volume was collected. For protein elution, four column volumes of elution buffer (TBS with either CaCl_2_ or EGTA as indicated in the figure) were added to elute proteins bound to the CaM-Agarose. Equivalent aliquots (15 µL) of each fraction were analysed by SDS-PAGE and immunoblotting using anti-His tag antibody (GenScript) overnight at 4°C as per manufacturer’s instructions. Immunoblots were visualized by chemiluminescence on a ChemiDoc Touch Imager (BioRad) and blots are representative of three replicate experiments.

### Dansyl-Calmodulin Binding Assays

CaM7 was dansylated as described previously (19). Briefly, 15 µL of 10 mM dansyl chloride in ethanol was added to 1 mL of 1 mg/mL CaM7 and was incubated at room temperature for 1 hour. The dansylated CaM7 (D-CaM) was then dialyzed against TBS overnight at 4°C. Steady-state D-CaM binding assays were performed by mixing 600 nM D-CaM with an equimolar amount of SUMO tag (negative control), SUMO-AtFBA4, or SUMO-AtFBA5 in either 1 mM CaCl_2_ or EGTA and incubated for 10 min at room temperature before reading in a SpectraMax Paradigm with 360 nm excitation and 400-600 nm emission settings. This experiment was performed in triplicate and graphed as the average trace of all replicates. D-CaM saturation binding assays were performed with the same settings as above except SUMO-AtFBA5 or SUMO-AtFBA5 peptide was added stepwise to 600 nM D-CaM and the emission maxima were determined from each replicate. The blue shift of the emission spectrum was used as a proxy for D-CaM saturation. This experiment was performed eight times and the saturation binding graph displays the mean peak wavelength at each SUMO-AtFBA5 concentration ± SD.

### Protein Sequence Alignments

Protein sequence alignments were performed using Clustal Omega (25) and visualized using Bioedit software (26).

### Statistical Analysis

All data was analyzed on GraphPad Prism 10. Statistical tests are indicated in figure legends.

## Results and Discussion

### Bioinformatic Analysis of Arabidopsis FBA Isozymes

Earlier studies indicate that the eight Arabidopsis FBA isozymes exhibit distinct localizations and functions (3, 4). Three isozymes, (FBA1, 2, 3) localize to the plastids and function predominantly in starch metabolism, while the remaining five isozymes are cytosolic and are involved in glycolytic processes (3, 6). FBA6 has also been detected in the nucleus (5). Previous work has shown the importance of AtFBA1 and AtFBA2 in carbon metabolism in the Calvin-Benson cycle and AtFBA3 in plastid respiration in non-photosynthetic tissue (4). In contrast, roles of AtFBA4 and AtFBA5 are less clear. A sequence comparison and phylogeny of the eight AtFBAs is presented in Figure 1A, B. Notable differences in their primary structures include the presence of an N-terminal plastid localization sequence in Arabidopsis FBA1, FBA2, and FBA3, as well as an N-terminal extension in FBA4, distinguishing it from the other cytosolic isozymes (Fig. 1). Interestingly, despite a high mitochondrial localization prediction score of the N-terminal extension of FBA4, co-localization and mitochondrial proteomic studies have demonstrated that AtFBA4 does not reside in the mitochondrion (3). However, some cytosolic FBAs have been found in association with the outer mitochondrial membrane, possibly as part of a glycolytic metabolon (27, 28). AtFBA4 and AtFBA5 are polypeptides of 393 and 358 residues, respectively, with predicted molecular masses of 42.9 and 38.3 kDa. Previous expression analysis (3) and an examination of transcriptomic databases using ePlant and Athena (29) suggest interesting differences in expression patterns among *FBA* genes (Fig. S1). *AtFBA4* is primarily expressed in floral tissue, especially pollen, but root expression also occurs, albeit at lower abundance relative to *FBA3*, *FBA6,* and *FBA8*. In contrast, *AtFBA5* is expressed exclusively in shoots and is significantly upregulated in response to cold, osmotic, drought, and salt stresses, as well as in the presence of exogenously applied sugars (3; Fig. S2). These distinct patterns suggest that AtFBA4 and AtFBA5 may play different roles under specific environmental conditions. In addition, phosphoproteomic studies identified several phosphorylation sites in both AtFBA4 and AtFBA5 (Fig. 1A) (PhosPhAt database; 30). Although, the effects of phosphorylation on AtFBA4 and AtFBA5 function are unknown, the control of glycolytic enzymes by reversible post-translational modifications (PTMs) is common (31, 32) and AtFBA6 is modified in response to cellular redox status (5).

**Figure 1.**
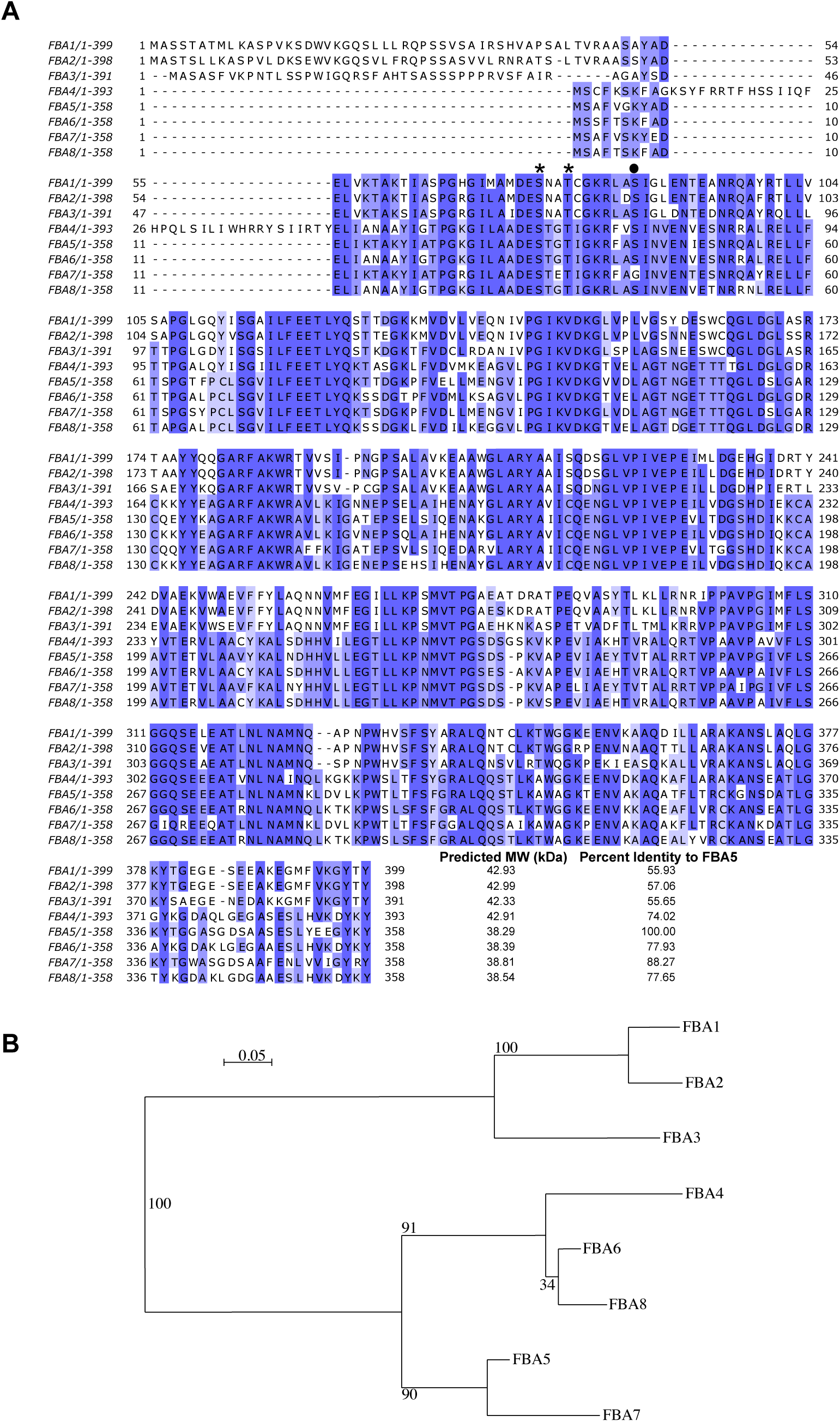
Comparison of Arabidopsis FBAs. (A) Multiple sequence alignment of Arabidopsis FBAs with 50% homology threshold for shading where darker shading indicates increasing level of similarity among the isoforms. Residue positions are indicated on the right. A solid circle above the sequences indicates a phosphorylation site identified in AtFBA5, whereas an asterisk indicates a phosphorylation site identified in both AtFBA4 and AtFBA5 (PhosPhAt database; 30). The predicted molecular weights (MW; kDa) of the AtFBAs and their respective percent identities to AtFBA5 are listed after the alignment. Locus identifiers for each AtFBA are as follows: *At2g21330* (AtFBA1), *At4g38970* (AtFBA2), *At2g01140* (AtFBA3), *At5g03690* (AtFBA4), *At4g26530* (AtFBA5), *At2g36460* (AtFBA6), *At4g26520* (AtFBA7), *At3g52930* (AtFBA8). (B) Maximum-liklihood phylogenetic tree of Arabidopsis FBAs constructed using PhyML as implemented in Seaview version 5.0.5 (51) with the muscle algorithm and the Blosum62 scoring matrix and other parameters at default values. Numbers at nodes correspond to percentage bootstrap frequencies for each node (100 iterations). The scale bar represents the average number of amino acid substitution per site.

### Characterization of recombinant AtFBA4 and AtFBA5

We expressed the AtFBA isozymes in *E. coli* with an N-terminally fused His-SUMO solubility tag, as the enzymes with a His tag alone were soluble but inactive after purification. Recombinant SUMO-AtFBA5 was purified to near homogeneity by Ni-affinity chromatography whereas SUMO-AtFBA4 copurified with several bacterial proteins (Fig. 2A), and further enrichment using size-exclusion chromatography resulted in an unstable enzyme unsuitable for analyses. Hence, we used preparations following Ni-affinity chromatography to preserve FBA activity, the caveat being that because AtFBA4 is only partially pure, its specific activity here is likely underestimated and we cannot exclude the possibility that the contaminating proteins may have affected its properties. Nevertheless, characterization of partially pure proteins is necessary when dealing with labile enzymes (22). Immunoblotting using anti-carrot Aldc immune serum (14) or anti-His antibodies confirmed the identity and intactness of recombinant SUMO-AtFBA4 at the expected molecular weight of 56 kDa, whereas none of the co-purifying bacterial proteins were immunoreactive (Fig. 2A). It is noteworthy that although AtFBA4 and AtFBA5 share 76% sequence identity, only AtFBA4 cross-reacted with our anti-Aldc immune serum, highlighting sequence differences between these isozymes.

**Figure 2.**
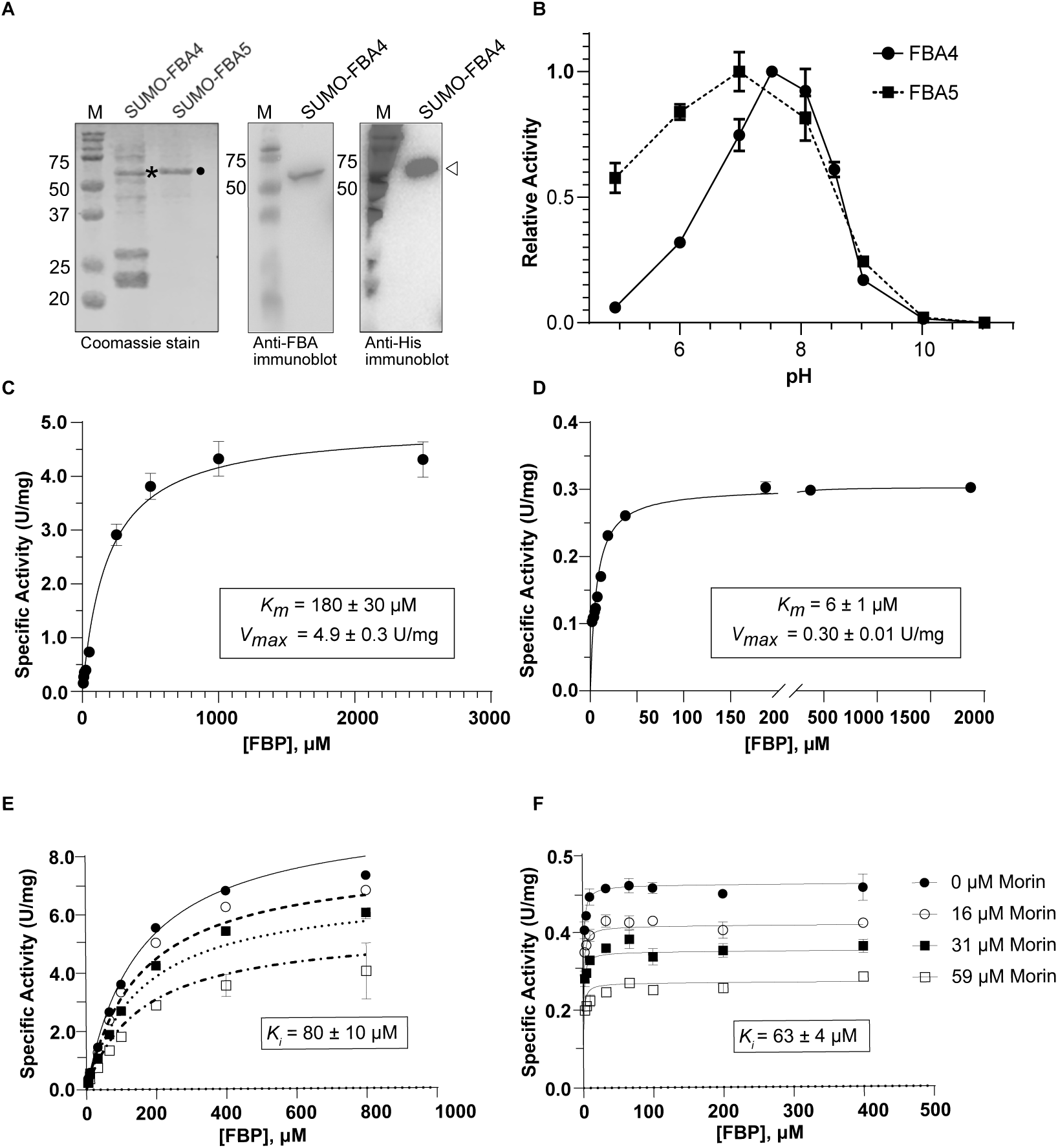
SDS-PAGE, immunoblot, and kinetic analysis of purified AtFBA4 and AtFBA5. (A) Coomassie stained SDS-PAGE (far left panel) of partially purified AtFBA4 (∼ 5 µg), and AtFBA5 (∼ 3 µg) as SUMO-fusion proteins used in subsequent assays. An asterisk and solid dot indicate the positions of SUMO-AtFBA4 and SUMO-AtFBA5, respectively. The positions of molecular weight markers (M, kDa) are indicated on the left. Immunoblots of recombinant SUMO-AtFBA4 following Ni-affinity purification. Following SDS-PAGE, samples were transferred to nitrocellulose and probed using anti-carrot FBA antiserum (middle panel) or anti-His antiserum (right panel) and detected using chemiluminescence. The immunoreactive SUMO-AtFBA4 is indicated by an arrowhead on the right. (B) Relative activity of SUMO-AtFBA4 and SUMO-AtFBA5 at different pH values assayed as outlined in the methods. Michaelis-Menten plots and the corresponding *K*_m_ and *V*_max_ values are shown for (C) SUMO-AtFBA4 and (D) SUMO-AtFBA5 for substrate fructose-1,6-bisphosphate (FBP). Inhibitory effects of morin, and the associated *K*_i_ values, are presented on FBP saturation curves of (E) SUMO-AtFBA4 and (F) SUMO-AtFBA5. All values represent the mean ±SD of *n* = 4 technical replicates.

Both isozymes exhibited optimal activity over a neutral pH range (7.0 - 7.5), which is typical for cytosolic FBAs (Fig. 2B) (14, 15). The apparent molecular masses of the isozymes using size-exclusion chromatography on a calibrated Superdex 200 Increase column were 240 ± 20 and 180 ± 20 kDa for AtFBA4 and AtFBA5, respectively (Fig. S3). This suggests homotetrameric complexes for both isozymes, consistent with previous reports on the quaternary structure of plant FBAs (14, 15).

With FBP as substrate, AtFBA4 and AtFBA5 displayed Michaelis-Menten kinetics with *V*_max_ and *K*_m_ values of 4.9 ± 0.3 U/mg and 180 ± 30 µM for AtFBA4, and 0.30 ± 0.01 U/mg and 6.0 ± 1 µM for AtFBA5, respectively (Fig. 2C, 2D). The specific activity for AtFBA5 was comparable with previous reports on Arabidopsis FBAs, whereas that of AtFBA4 was considerably higher (5, 6). As noted above, our specific activity estimate for AtFBA4 is likely an underestimation. The *K*_m_(FBP) value of AtFBA4 was similar to previous studies on Arabidopsis FBAs, whereas AtFBA5 displays much stronger FBP affinity, with only a cytosolic FBA from germinating castor oil seeds exhibiting a lower *K*_m_(FBP) value of 0.16 µM (15). These *K_m_* values are consistent with physiological concentrations of FBP reported for Arabidopsis rosettes (33).

### Inhibitory Effect of Morin on AtFBA4 and AtFBA5 Activity

The flavonol morin is a specialized plant metabolite and potent inhibitor of protozoan FBAs, and there is considerable interest in using morin as a potential therapeutic agent (17, 18). Thus, we investigated its effects on AtFBA4 and AtFBA5 activity. Similar to its action on EtFBA2 (17), morin exerted non-competitive inhibition of AtFBA4 and AtFBA5, with *K*_i_ values of 80 ±10 and 63 ±4 µM for AtFBA4 and AtFBA5, respectively (Fig. 2E, F). Arabidopsis produces morin-related flavonoids such as quercetin and derivatives, some of which have been implicated in development and stress-response pathways (34, 35). The *K*_i_ values we observed for morin would be expected to lie within the µM range of flavonoids that are induced during biotic and abiotic stress (36). Whether glycolytic activity in plants is affected by morin-related flavonoids remains an open question, but it is worth noting that both cytosolic pyruvate kinase and phosphoenolpyruvate carboxylase from *Brassica napus* cell cultures are potently inhibited by the flavonoids rutin and quercetin (37, 38). Future studies should explore whether the anti-bacterial and anti-tumor properties attributed to morin and related flavonoids may be due to their ability to reduce glycolytic flux by inhibiting bacterial and human FBA isozymes. Given the structural similarities between morin and quercetin, it may be productive to test flavonoid libraries in the future for novel inhibitors of plant FBAs and explore their roles *in vivo*. We note that none of the other metabolites we tested inhibited AtFBA4 or AtFBA5 activity to the degree of morin (Table S2).

### CaM Interacts with AtFBA5 in a Ca^2+^-dependent manner

A previous study suggested that *M. sativa* MsFBA6 interacts with CaM-like 10 protein, MsCML10 (8). To determine whether this property is conserved with the Arabidopsis orthologs, we used *in planta* split-luciferase (SL) assays to test the interaction between AtFBA4 and AtCML11, the closest respective Arabidopsis orthologs of MsFBA6 and MsCML10. We selected AtFBA5 as a negative FBA control for interaction with AtCML11, and used CaM7, AtCML8, and AtCML42 as additional negative controls based on their high homology to AtCML11 and our previous study showing strong expression of these proteins in the SL system (20). We did not observe interaction of AtFBA4 with any of the CaM/CMLs tested in the SL system (Fig. 3). Surprisingly, AtFBA5 exhibited a strong and specific interaction with CaM7, an evolutionarily-conserved CaM isoform (Fig. 3A, B). Direct binding between CaM7 and AtFBA5 was further confirmed through two independent *in vitro* assays: Ca^2+^-dependent CaM-agarose binding (Fig. 3C) and steady-state dansyl-CaM7 fluorescence (Fig. 3D, E). CaM7 interacted specifically with AtFBA5 in a Ca^2+^-dependent manner, but not with AtFBA4 or the SUMO tag alone. Using dansylated-CaM7 (D-CaM), we estimated binding affinity (*K*_D_) of the Ca^2+^-CaM/AtFBA5 interaction at 190 ± 40 nM (Fig. 3F), which falls within the physiological range for CaM interactors (39, 40).

**Figure 3.**
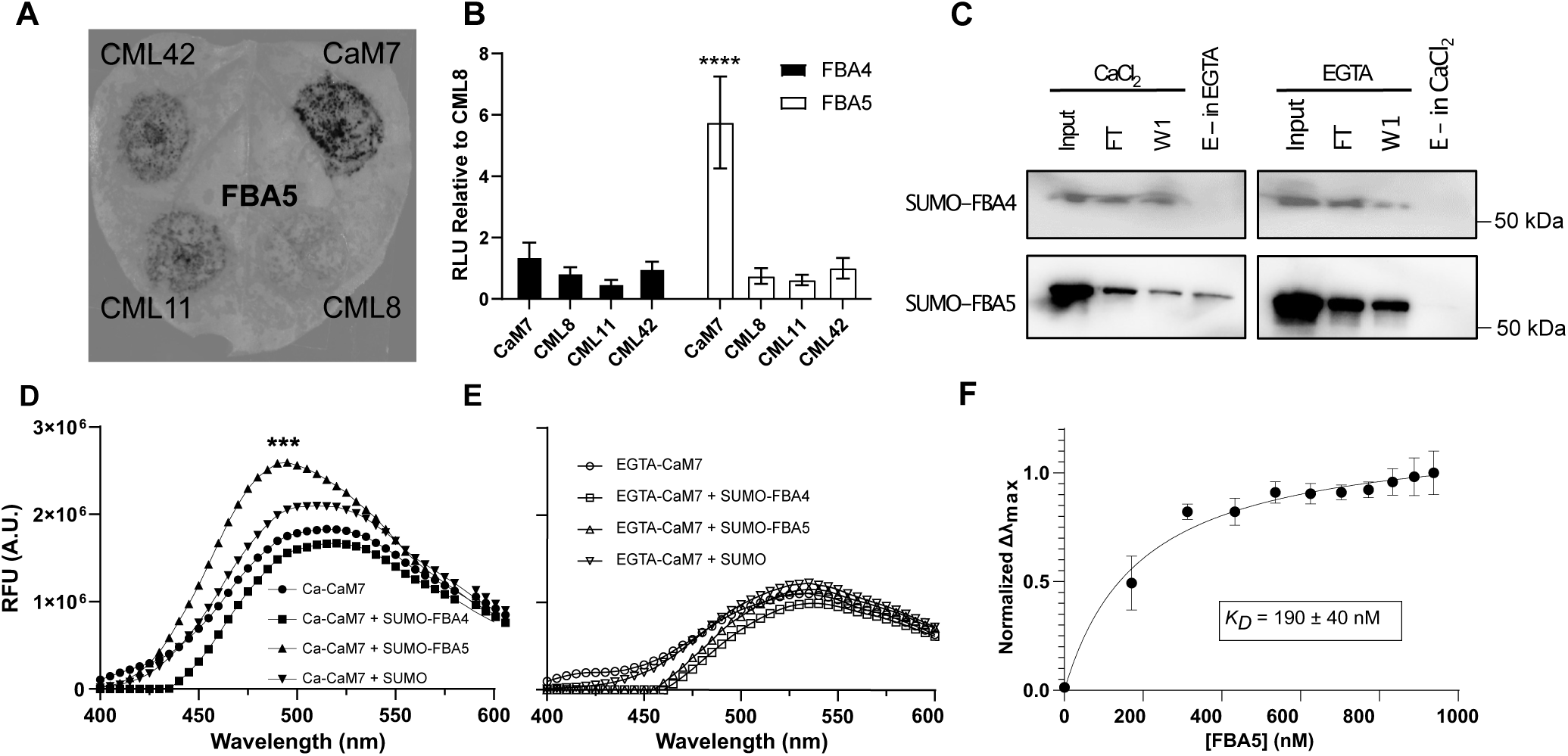
FBA5 specifically interacts with calmodulin (CaM) in a Ca^2+^-dependent manner. The split-luciferase system was used to test the interaction of fusion protein NLuc-AtFBA5 or NLuc-AtFBA4 with CLuc-fusions of -CaM7 (a conserved CaM isoform), -AtCML8, - AtCML11, or -AtCML42 *in planta* using *N. benthamiana* leaves as described in Methods. (A) A qualitative, representative whole-leaf image is shown for split-luciferase assays of NLuc-AtFBA5 with the different CLuc-CaM/CMLs. (B) Bar graphs showing quantitative analysis (representative of 4 biological replicates) of split-luciferase assays testing interaction of NLuc-AtFBA4 and -AtFBA5 with CaM/CMLs *in planta*. Data are expressed relative to the RLU (relative light unit) signal observed using the negative control AtCML42 (set to an RLU = 1.0). (C) CaM-affinity chromatography was used to assess the *in vitro* binding of recombinant SUMO-AtFBA4 or -AtFBA5 to CaM-sepharose in the presence of 1 mM CaCl_2_ or 5 mM EGTA as indicated. Anti-His immunoblots are presented where input, *FT*, *W1*, and *E* indicate protein input, flow-through (unbound eluate), column wash, and eluted sample, respectively. (D) *In vitro* binding of CaM to SUMO-AtFBA4 and -AtFBA5 was also tested using 600 nM dansylated-CaM7 and observing changes in dansyl fluorescence in the presence of (D) 1 mM CaCl_2_ or (E) 5 mM EGTA. Panels *D* and *E* share the same vertical axis. Average fluorescent-spectra are shown for dansyl-CaM7 incubated with SUMO (negative control), SUMO-AtFBA4, or SUMO-AtFBA5. (F) Binding of CaM to AtFBA5 was monitored by the shift in peak fluorescence wavelength of SUMO-AtFBA5 titrated into 600 nM dansylated CaM7 in the presence of 1 mM CaCl_2_. The saturation plot shown is representative of three replicates where data points and the *K*_D_ are the mean ± SD of eight technical replicates. All other quantitative data are the mean values or mean ± SD where applicable (One-way ANOVA against AtCML8 (B) or CaM7 alone (D, E) with Sidak’s test for multiple comparisons, p-value * < 0.05, ** < 0.01, *** < 0.001, **** < 0.0001).

There are several possible explanations as to why we did not observe binding of AtFBA4 to AtCML11 as reported for their *Medicago* orthologs (8). Our *in planta* analysis conditions may not have been suitable for the interaction, or this association may not be conserved across plant taxa. Indeed, it is noteworthy that at 79 % identity, MsCML10 and AtCML11 are less conserved as orthologs than are MsFBA6 and AtFBA4 (86 % identity). Moreover, AtCML11 possesses an unusual N-terminal poly-Q extension which is not present in MsCML10 (Fig. S4). These structural differences between the CML orthologs may explain the lack of interaction between AtCML11 and AtFBA4 compared to the *Medicago* orthologs.

### CaM binding domain identification

To delineate the CaM-binding domain (CaMBD) within AtFBA5, we tested CaM7 binding to a series of AtFBA5 truncated proteins using the SL system (Fig. 4A). AtCML8 was used to normalize the interaction signal as it is among the CMLs with the highest homology (73% identity) to conserved CaM (12). AtFBA5 amino acid (AA) residues 180-270 exhibited the strongest interaction with CaM7 compared to controls. Within this 90 AA sequence, a single CaMBD of approximately 30 residues was predicted using an online algorithm (41). This putative CaMBD associated with CaM7 in a Ca^2+^-dependent manner as a SUMO-fusion protein yielding a *K*_D_ of 290 ± 130 nM (Fig. 4B). This *K*_D_ is higher, but within error of the value (190 ± 40 nM) observed using full-length AtFBA5 (Fig. 3F), indicating that the full-length FBA5/CaM complex may be more stable than the CaMBD-peptide/CaM complex. We note, however, that although the full-length AtFBA5 interacted with CaM7 in our SL and *in vitro* assays (Fig. 3), as did the truncated region 180-270, several other fragments which included the putative CaMBD (∼190-220) did not show interaction in SL assays (Fig. 4A). The reason for this is unclear but may have been due to poor expression or instability of these partial proteins when expressed transiently in *N. benthamiana* SL assays. Nevertheless, the high-affinity, Ca^2+^-dependent interaction between CaM7 and full-length AtFBA5 suggests that this binding may occur during stresses when cytosolic Ca^2+^ concentrations become elevated. The CaMBD of AtFBA5 most closely resembles a 1-5-10 CaM-binding motif (Fig. 4C), with a predicted amphiphilic alpha-helical structure (Fig. 4D) carrying a net positive charge, characteristics that are typical of CaMBDs (9, 40). Interestingly, in comparing this region in AtFBA4 and AtFBA5, there are several non-conserved changes, including a K to E substitution in AtFBA4 (Fig. 4C). These differences may account for the inability of AtFBA4 to bind CaM7, but this remains speculative.

**Figure 4.**
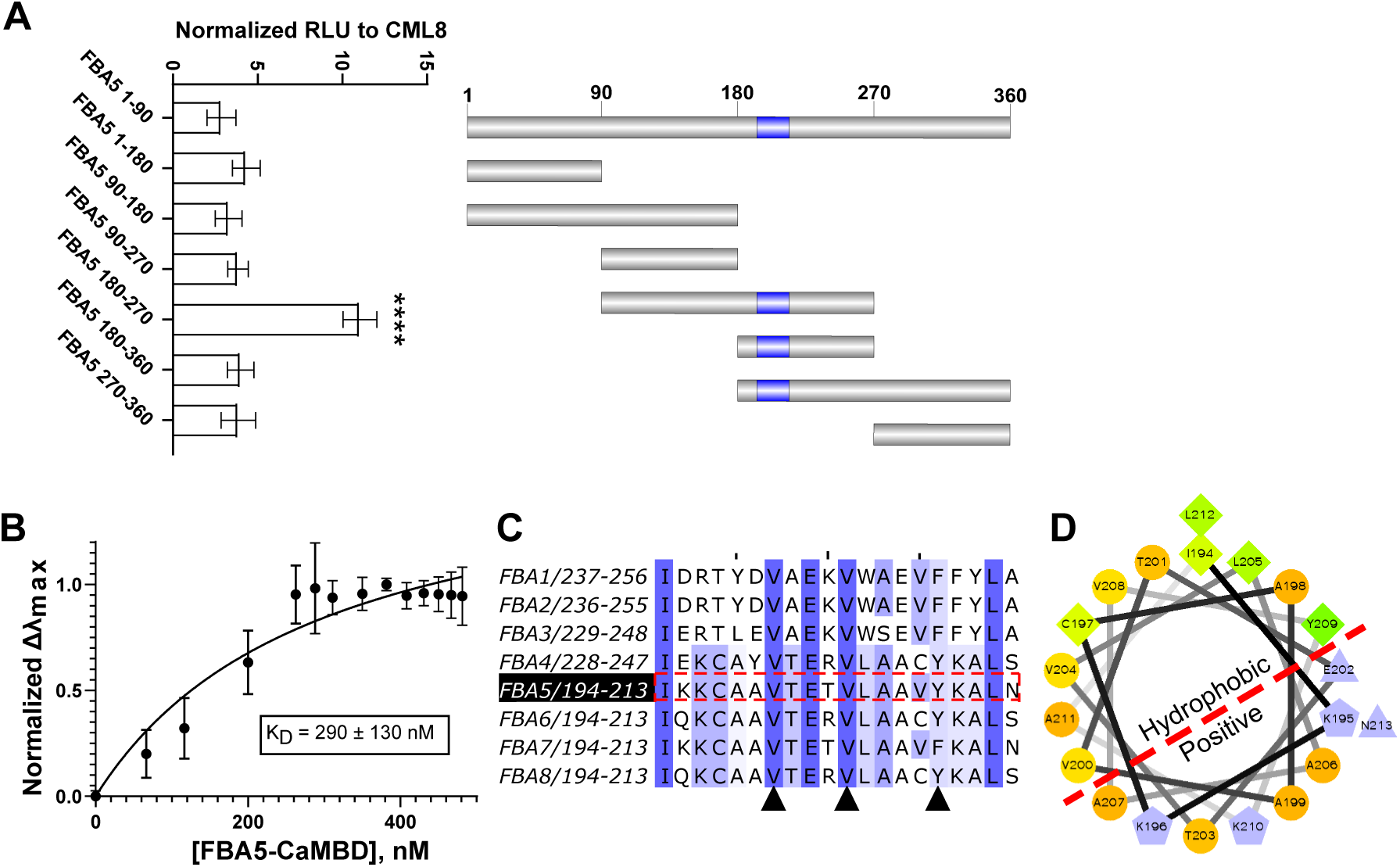
Identification of the putative AtFBA5 CaM binding domain (CaMBD). (A) Quantitative representative split-luciferase assays of NLuc-AtFBA5 delineations tested for their interaction with CLuc-CaM7 and plotted as RLU normalized to CLuc-AtCML8. The NLuc-AtFBA5 delineations tested are graphically represented in the right-hand panel with the relative position of the CaM binding domain colored in purple. The top schematic image represents the full-length AtFBA5 protein where numbers indicate residue positions. (B) Saturation binding curve of SUMO-AtFBA5 CaMBD (residues 194-213) titrated into 600 nM dansylated Ca^2+^-CaM7 and monitored by the shift in peak fluorescence wavelength. (C) Multiple sequence alignment of the putative AtFBA5 CaMBD (residues 194-213) with the corresponding regions in the Arabidopsis FBA paralogs. Arrows represent possible hydrophobic contact residues in the CaMBD of AtFBA5. The sequence is shaded based on similarity where dark shading is 100% identity, lighter shading is similarity, and no shading is less than 50% similarity among isoforms. (D) Helical wheel diagram of the putative AtFBA5 CaMBD showing the hydrophobic and positively charged faces of the alpha-helix that is typical for CaMBDs. Single-site saturation binding kinetics (panels A and C) were determined using GraphPad Prism 10, and a One-way ANOVA with Sidak’s test for multiple comparisons was used to assess statistical significance (panel B), p-value * < 0.05, ** < 0.01, *** < 0.001, **** < 0.0001).

### Ca^2+^-CaM binding increases stability of AtFBA5

In a previous study, overexpression of MsCML10 in *M. sativa* led to increased aldolase activity in cell extracts, indirectly supporting the hypothesis that MsCML10 activates MsFBA6 activity (8). However, we did not observe binding of CaM7 or AtCML11 to AtFBA4, the closest ortholog of MsFBA6. Although AtFBA5 binds to CaM7, under our *in vitro* assay conditions, we did not observe any effect of CaM7 on the catalytic activity of AtFBA5 (Fig. S5). We then examined the impact of CaM7 on the stability of AtFBA5 across a range of temperatures (Fig. 5). In the absence of Ca^2+^, AtFBA5 and the CaM7-AtFBA5 mixture exhibited a comparable T_1/2_ of 45-46 ± 2°C. However, when Ca^2+^ was present, the CaM7-AtFBA5 complex demonstrated a statistically significant increase in melting temperature, with an estimated T_1/2_ of 50 ± 2°C (Fig. 5A). This ∼5°C increase in thermal stability is similar in magnitude to that reported for other proteins as a result of protein-protein interaction (42, 43).

**Figure 5.**
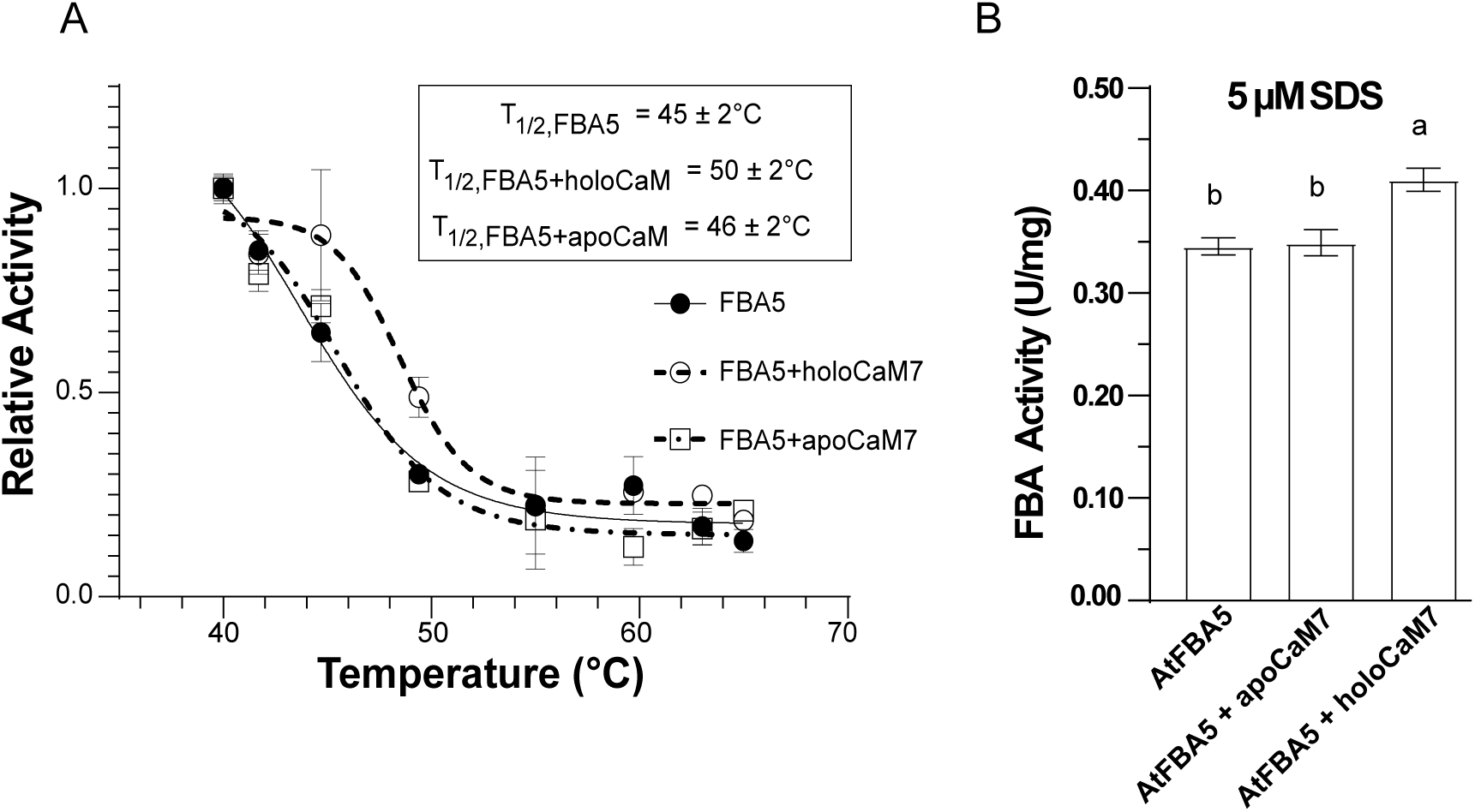
Ca^2+^-CaM binding to AtFBA5 increases its thermal and chemical stability. (A) SUMO-AtFBA5 (∼1 mg/mL) was aliquoted into PCR tubes without CaM (FBA5) or with CaM in molar excess with 1 mM CaCl_2_ (holoCaM) or 1 mM EGTA (apoCaM) and incubated for 20 min in a thermocycler over a range of temperatures followed by FBA activity assays as described in Methods. Relative FBA activity was plotted against the incubation temperature and the T_1/2_, defined as the temperature at which the enzyme retains 50% of its activity after the 20 min heat treatment, was calculated from the sigmoidal curve using GraphPad Prism 10. All points represent the mean relative T_1/2_ activity (± SD) of *n* = 4 technical replicates. (B) SUMO-AtFBA5 *in vitro* activity was measured in the presence of 5 µM SDS. Samples, as indicated included SUMO-AtFBA5 alone (AtFBA5), or in the presence of 200 nM CaM7 and 1 mM EGTA (AtFBA5 + apoCaM7), or with 200 nM CaM7 and 1 mM CaCl_2_ (AtFBA5 + holoCaM7).

At low concentrations, the detergent SDS is often used as a chemical denaturant to assess protein stability (44, 45). Thus, we tested the impact of 5 µM SDS on AtFBA5 activity in the absence or presence of CaM7 (Fig. 5B). Similar to our results in heat-stability tests, AtFBA5 activity was greater in the presence of SDS only when both Ca^2+^ and CaM7 were included in the assays. These data further provide further support for CaM7-AtFBA5 interaction and indicate that CaM7 stabilizes the AtFBA5 enzyme in a Ca^2+^-dependent manner. The stabilization of AtFBA5 by CaM7 is reminiscent of a previous study on CTP:phosphocholine cytidylyltransferase where CaM binding stabilizes and protects the enzyme from proteolysis but does not alter its kinetic properties (46), and a study showing CaM-mediated stabilization of erythrocyte protein 4.1R (47).

## Concluding Remarks

Our data indicate that AtFBA4 and AtFBA5 function as homotetramers with high FBP affinity and properties that are consistent with their predicted roles as cytosolic FBAs. It is also noteworthy that AtFBA5 displayed the lowest *K_m_*(FBP) value described for any Arabidopsis FBA to date. The interaction reported for MsFBA6 and MsCML10 (8) is not conserved in the closest Arabidopsis orthologs AtFBA4 and AtCML11 but, interestingly, AtFBA5 is stabilized by specific, high-affinity CaM binding in a Ca^2+^-dependent manner. Given that *AtFBA5* is upregulated by a variety of abiotic stresses (Fig. S2), we speculate that Ca^2+^-CaM binding to AtFBA5 helps to stabilize the enzyme when cells are challenged with stressed conditions and Ca^2+^ levels are elevated. Moreover, stabilization of the AtFBA5 complex may also provide a means to integrate stress signaling with sucrose metabolism. The latter process is tightly linked to stress response given that plants undergo metabolic reprogramming and osmoprotectant synthesis to cope with adverse environmental conditions (48, 49).

Similar to a recent report on the inhibition of *E. tenella* FBAs by the plant flavonoid morin (17), we found morin to be a potent inhibitor of AtFBA4 and AtFBA5, raising questions about a potential regulatory role for flavonoids in plant glycolysis. Flavonoids participate in diverse physiological processes, including defense against UV radiation, pathogen attacks, low nutrient, and oxidative stress (48, 36). Interestingly, a previous study found that the addition of flavonoids to growth media inhibits germination and plant growth (50), but whether the underlying mechanisms involve inhibition of glycolytic flux remain unclear. Under stress conditions, the modulation of FBA activity by stress-responsive factors, such as flavonoids and Ca^2+^, may serve to regulate carbon flow and energy utilization.

## Abbreviations

FBA: fructose bisphosphate aldolase
FBP: fructose 1,6-bisphosphate
CaM: calmodulin
CaMBD: CaM-binding domain
CML: calmodulin-like
DHAP: dihydroxyacetone phosphate
G3P: glyceraldehyde-3-phosphate
SL: split-luciferase
SUMO: small ubiquitin-like modifier
TBS: Tris-buffered saline
U: unit of FBA activity

## Supporting Data

**Supporting Table S1:**
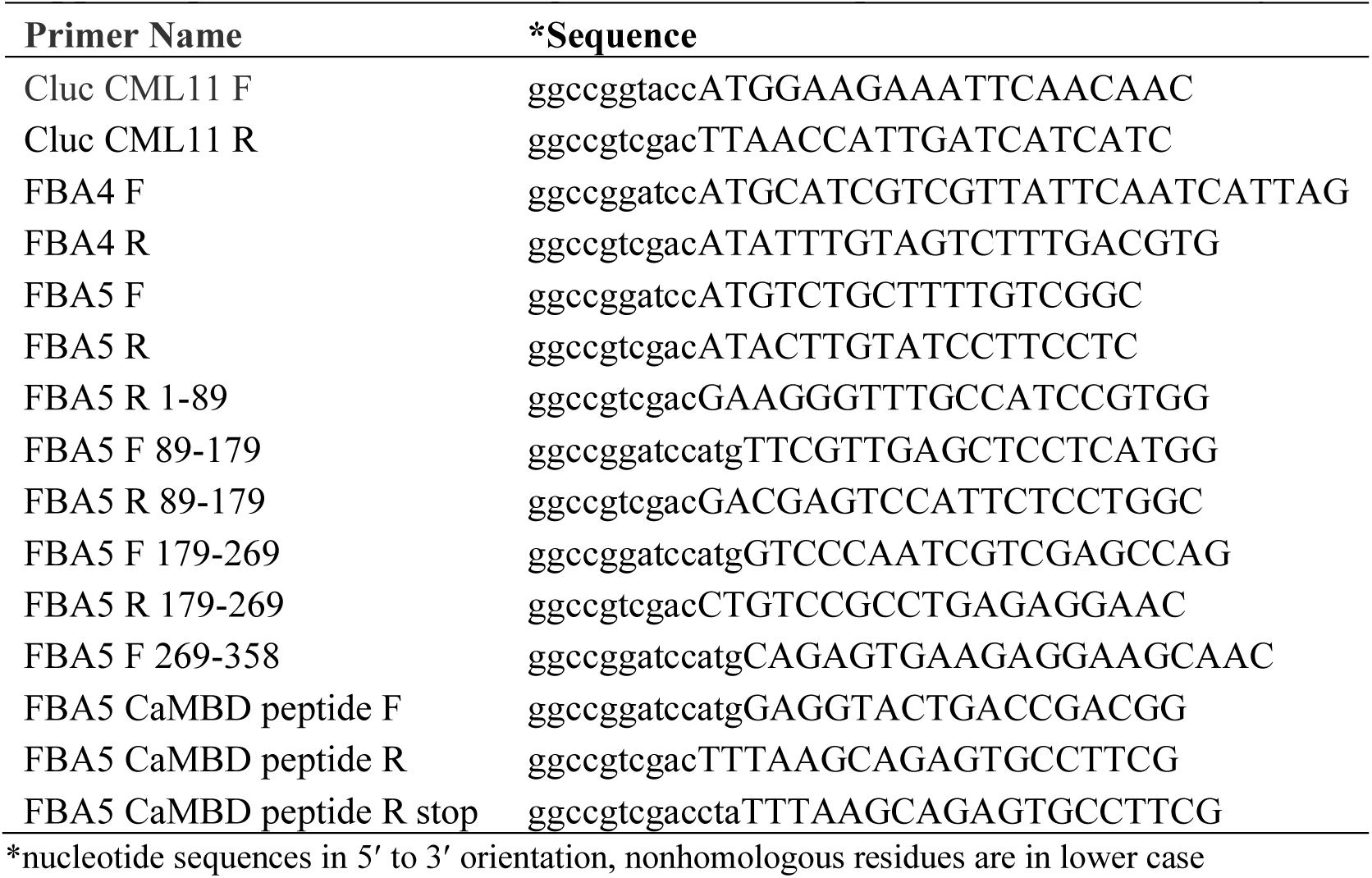
List of oligonucleotide PCR primers used in this study.

**Supporting Table S2.** Influence of various metabolites on FBA4 and FBA5 activity. All assays were conducted at pH 7.5 with 250 μM FBP. Enzymatic activity in the presence of effectors is expressed as mean ± SD relative to the respective control set at 100%. Data represent a minimum of two biological replicates with a minimum of three technical replicates each.

|  | Concentration<br>(mM) | FBA4 |  | FBA5 |  |
| --- | --- | --- | --- | --- | --- |
|  |  | Mean | SD | Mean | SD |
| Control | - | 100 | 3 | 100 | 6 |
| R5P | 1.0 | 100 | 3 | 111 | 5 |
| G6P | 1.0 | 102 | 5 | 113 | 7 |
| F-2,6-P | 0.20 | 101 | 5 | 117 | 4 |
| T6P | 1.0 | 103 | 3 | 116 | 3 |
| Malate | 5.0 | 68 | 12 | 107 | 1 |
| Succinate | 5.0 | 89 | 5 | 98 | 6 |
| 2-OG | 5.0 | 69 | 2 | 102 | 2 |
| ACoA | 1.0 | 88 | 4 | 88 | 4 |
| PEP | 1.0 | 85 | 6 | 102 | 17 |
| Citrate | 5.0 | 89 | 5 | 109 | 6 |
| Asp | 3.0 | 91 | 3 | 93 | 7 |
| Glu | 5.0 | 102 | 2 | 99 | 8 |
| Tyr-Asp | 0.50 | 100 | 2 | 82 | 5 |
| GSSG | 0.75 | 104 | 3 | 80 | 11 |
| PP<sub>i</sub> | 1.0 | 87 | 2 | 144 | 10 |
| P<sub>i</sub> | 5.0 | 94 | 2 | 101 | 7 |
| ATP | 1.0 | 85 | 1 | 87 | 5 |
| AMP | 1.0 | 101 | 2 | 84 | 8 |
| Morin | 0.065 | 67 | 6 | 12 | 2 |

**Supporting Data Figure S1.**
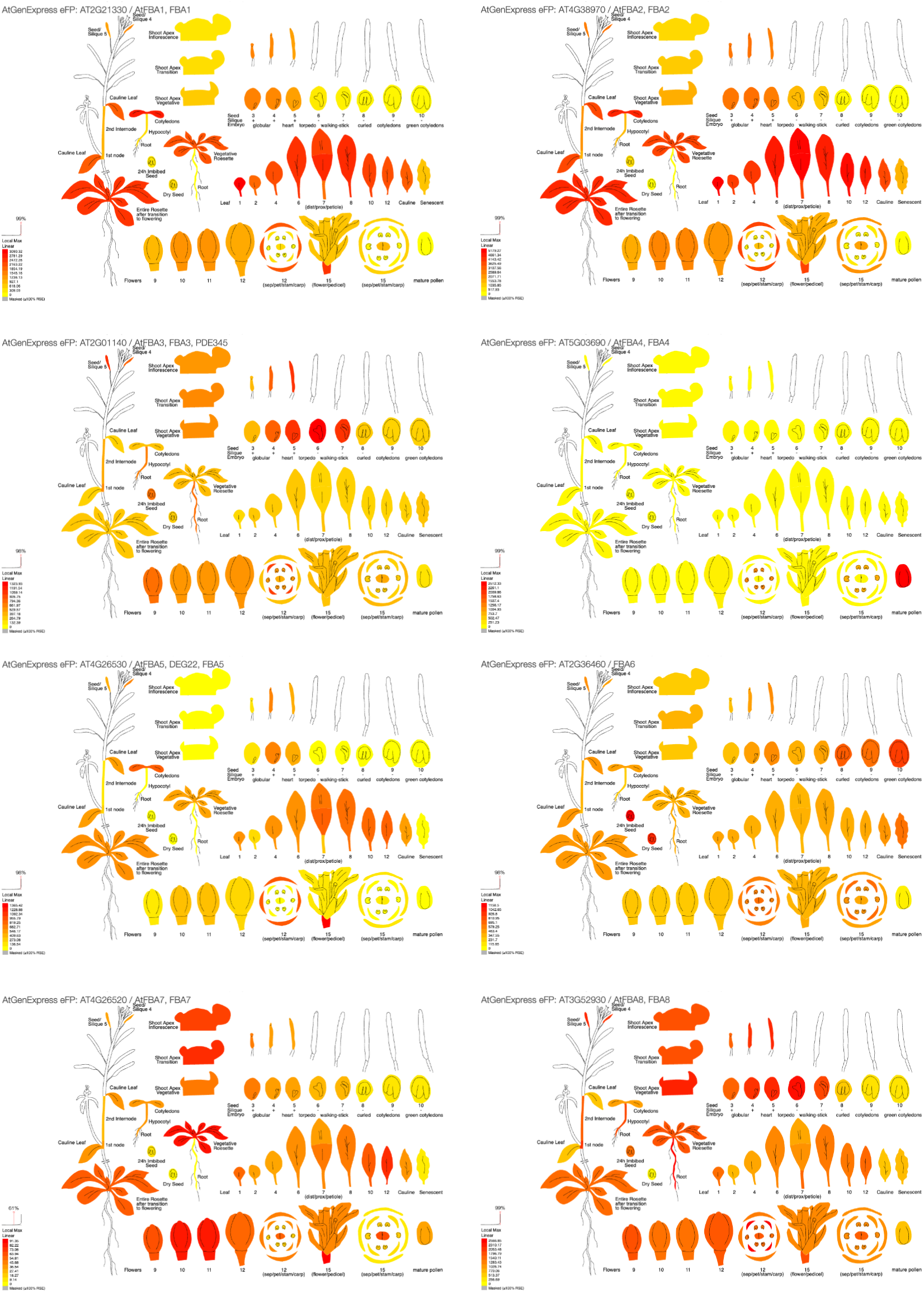
Developmental and tissue expression profiles of the eight Arabidopsis *AtFBA* aldolase genes. Data and images were derived from the ePlant visualization tool in the online Bio-Analytical Resource for Plant Biology (BAR, https://bar.utoronto.ca/eplant/).

**Supporting Data Figure S2.**
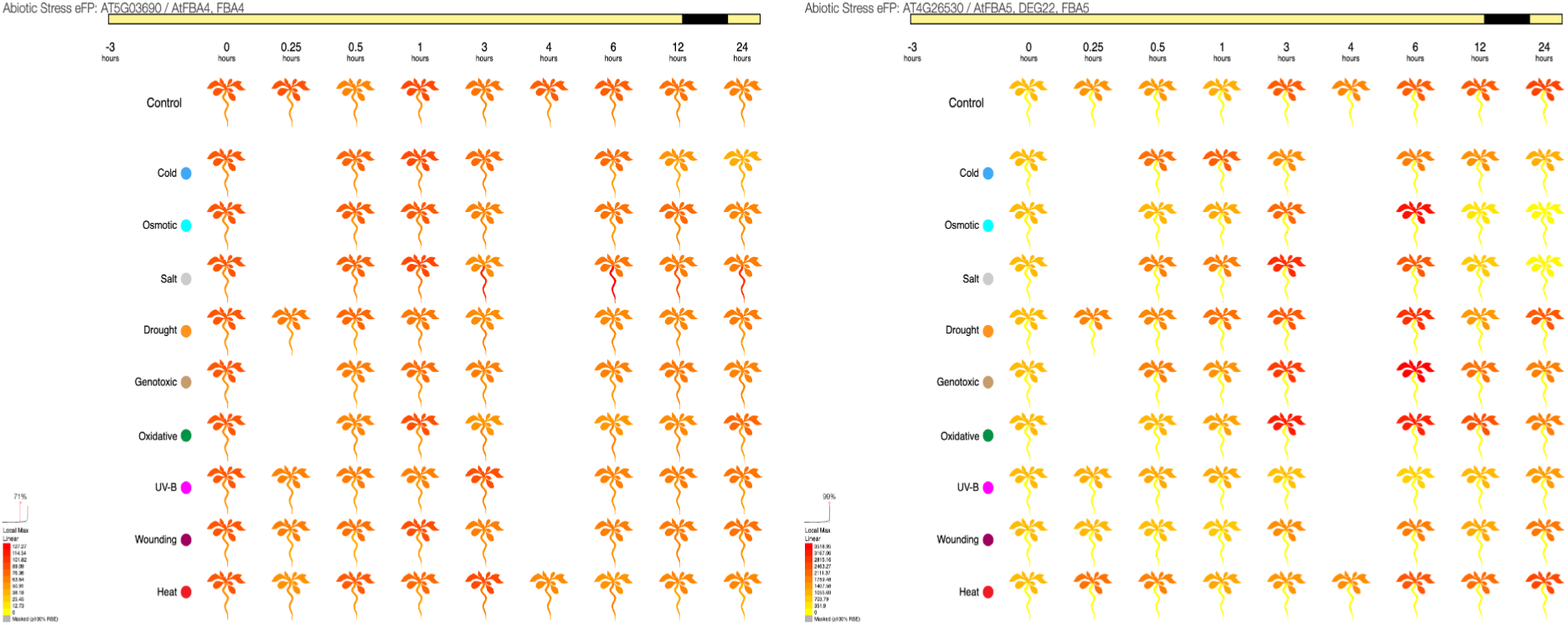
Gene expression profiles of Arabidopsis *AtFBA4* and *AtFBA5* in response to various abiotic stress treatments. Data and images were derived from the ePlant visualization tool in the online Bio-Analytical Resource for Plant Biology (BAR, https://bar.utoronto.ca/eplant/).

**Supporting Data Figure S3:**
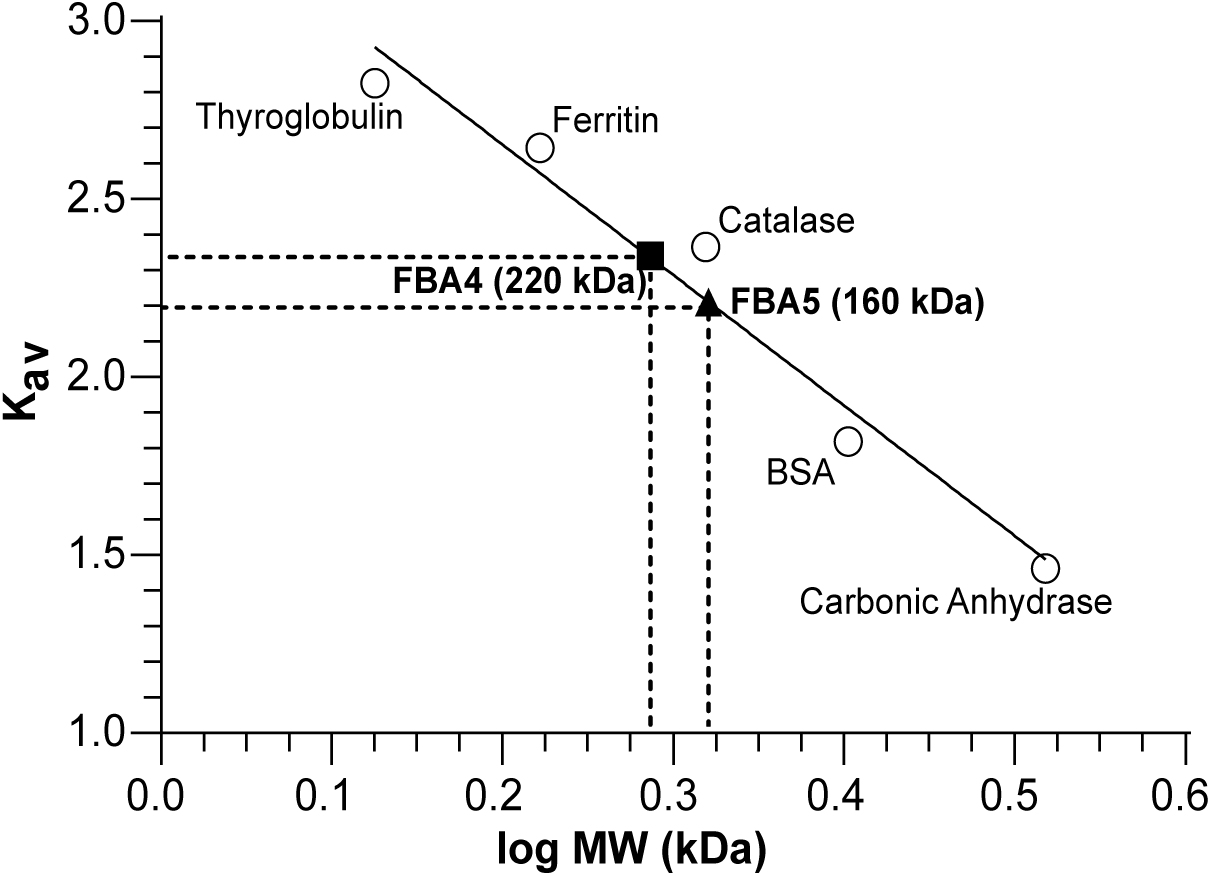
Estimation of molecular masses of aldolases AtFBA4 and AtFBA5. Size-exclusion chromatography, using various protein standards as indicated on the figure, was used to generate a calibration curve to estimate the native molecular masses of SUMO-FBA4 and SUMO-FBA5 as described in Methods.

**Supporting Data Figure S4.**
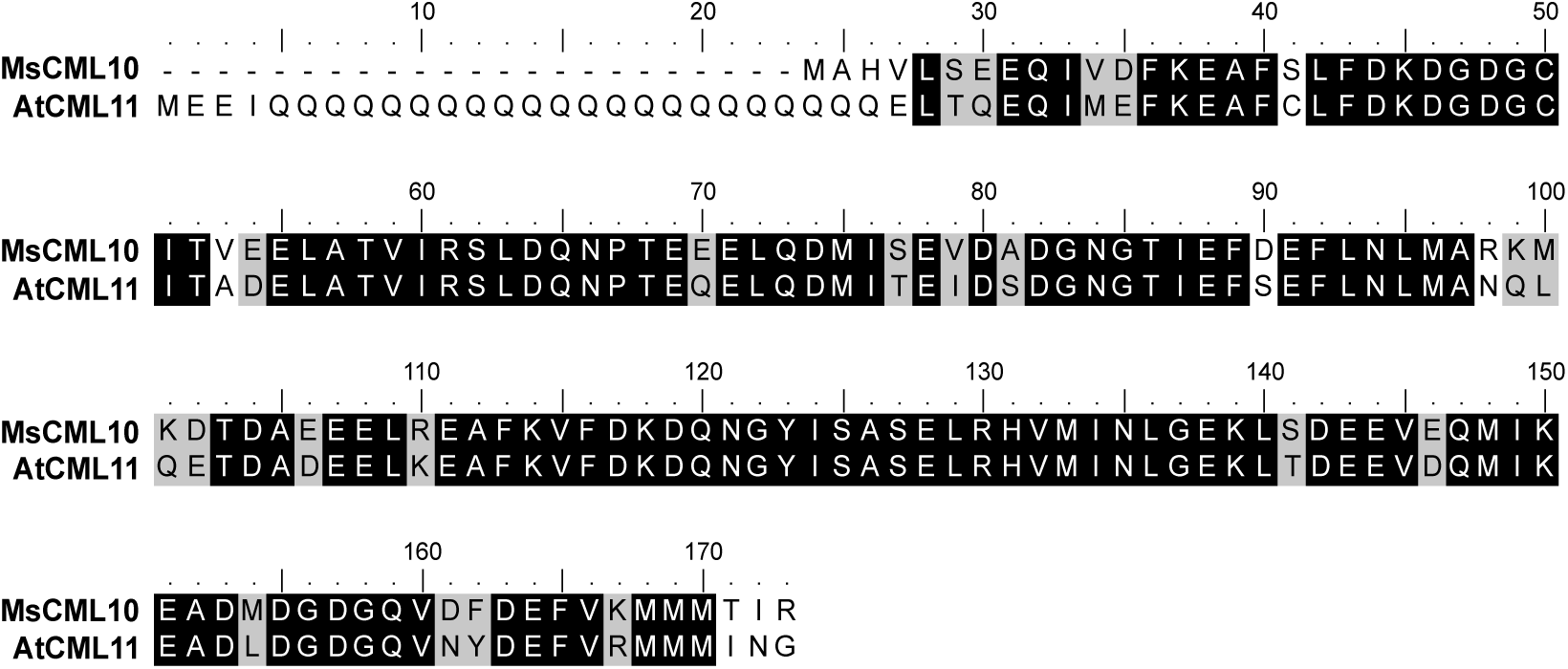
Protein sequence alignment of *Medicago sativa* CML10 and *Arabidopsis thaliana* CML11. Similar and identical residues are shaded in grey and black, respectively. Alignment was performed using Clustal Omega (Sievers and Higgins, 2014) and the shaded-box image created in Bioedit (Hall, 1999).

**Supporting Data Figure S5.**
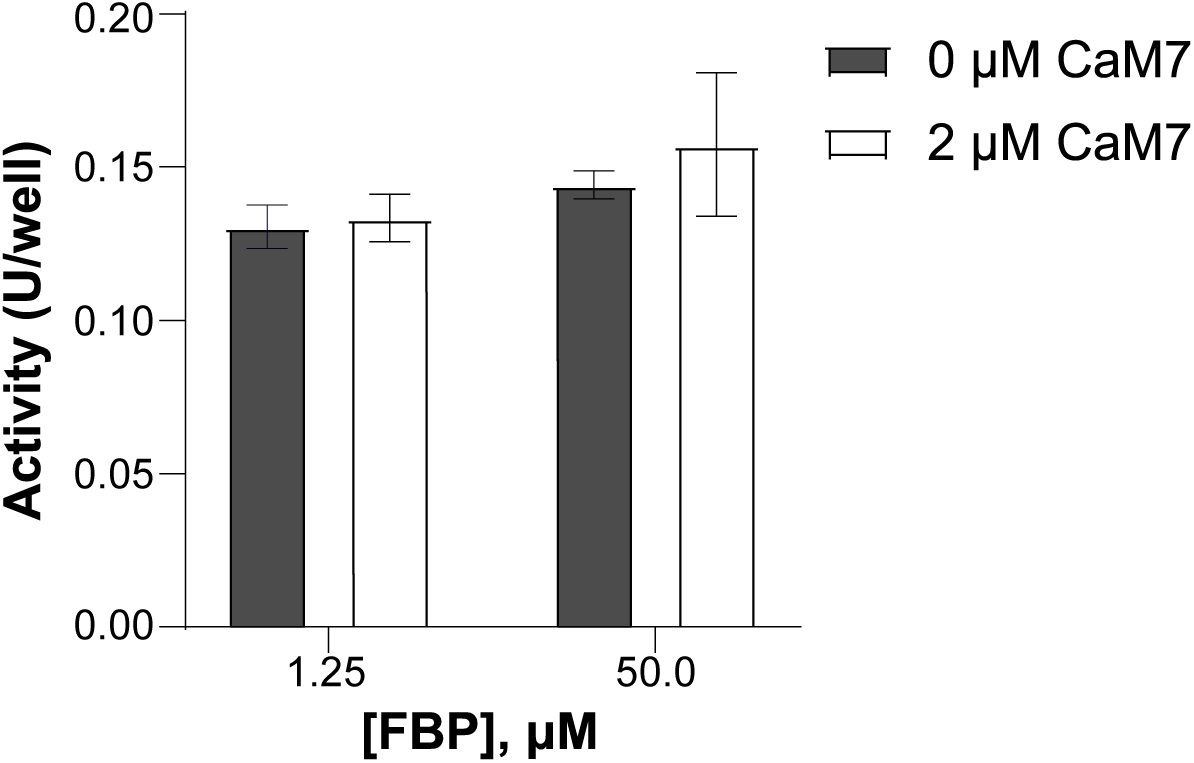
CaM does not alter *in vitro* AtFBA5 activity. SUMO-AtFBA5 activity was assayed, at the substrate concentrations indicated, as described in Methods in either the absence or presence of CaM. For assays with CaM, CaCl_2_ was included at 1 mM.

**Supporting Data Figure S6.**
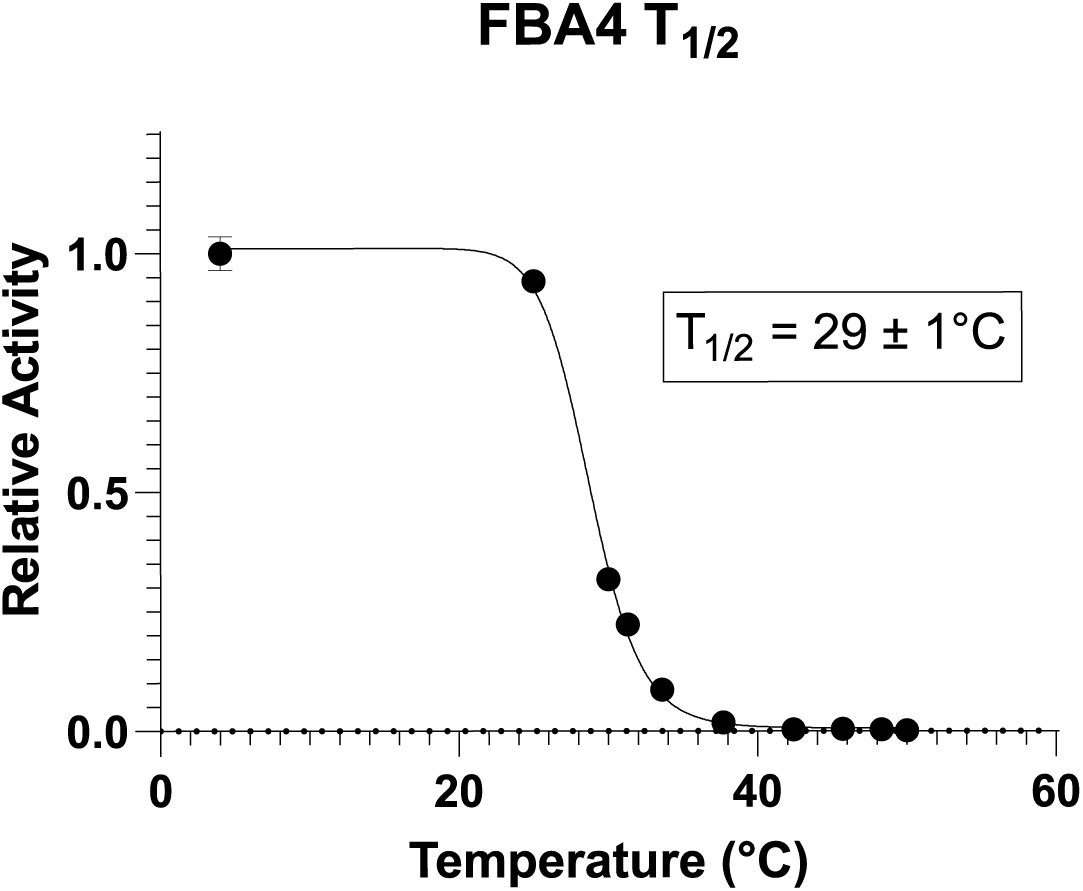
SUMO-AtFBA4 (∼1 mg/mL) was aliquoted into PCR tubes and incubated for 20 mins in a thermocycler over a range of temperatures followed by FBA activity assays as described in Methods. Relative FBA activity was plotted against the incubation temperature and the T_1/2_, defined as the temperature at which the enzyme retains 50% of its activity after the 20 min heat treatment, was calculated from the sigmoidal curve using GraphPad Prism 10. All points represent the mean relative T_1/2_ activity (± SD) of *n* = 4 technical replicates.

